# Neurotrophin signaling is modulated by specific transmembrane domain interactions

**DOI:** 10.1101/2021.05.24.445441

**Authors:** María L. Franco, Kirill D. Nadezhdin, Taylor P. Light, Sergey A. Goncharuk, Andrea Soler-Lopez, Fozia Ahmed, Konstantin S. Mineev, Kalina Hristova, Alexander S. Arseniev, Marçal Vilar

**Affiliations:** Unit of Molecular Basis of Neurodegeneration, Institute of Biomedicine CSIC. 46010 València, SPAIN; Shemyakin-Ovchinnikov Institute of Bioorganic Chemistry of the Russian Academy of Sciences, Moscow 117997, Russian Federation; Department of Materials Science and Engineering, Johns Hopkins University, Baltimore, MD, USA

## Abstract

The neurotrophin receptors p75 and TrkA play an important role in the development and survival of the nervous system. Biochemical data suggest that p75 and TrkA regulate the activities of each other. For instance, p75 is able to regulate the response of TrkA to lower concentrations of NGF and TrkA promotes p75 shedding by α-secretases in a ligand-dependent manner. The current model is that p75 and TrkA are regulated by means of a physical direct interaction, however the nature of such interaction has been elusive so far. Here using NMR in micelles, multiscale molecular dynamics (MD), FRET and functional studies we identified and characterized the direct interaction between TrkA and p75 through the transmembrane domains (TMDs). MD of p75-TMD mutants suggests that although the interaction between TrkA and p75 TMDs is maintained, a specific protein interface is required to facilitate TrkA active homodimerization in the presence of NGF. The same mutations in the TMD protein interface of p75 reduced the activation of TrkA by NGF and cell differentiation. In summary we provide a structural model of the p75/TrkA receptor complex stabilized by transmembrane domain interactions.

## Introduction

Nerve growth factor (NGF) is a member of the mammalian neurotrophin (NT) protein family, which also includes BDNF, NT3, and NT4/5 (1). NTs are implicated in the maintenance and survival of the peripheral and central nervous systems and mediate several forms of synaptic plasticity (2–5). NTs interact with two distinct receptors, a cognate member of the Trk receptor tyrosine kinase family and the common p75 neurotrophin receptor, which belongs to the tumor necrosis factor receptor (TNFR) superfamily of death receptors (6, 7). While Trk receptor signaling is involved in survival and differentiation (8, 9), p75 participates in several signaling pathways (reviewed in (10)). p75-mediated signaling is governed by the cell context and the formation of complexes with different co-receptors and ligands, such as sortilin/pro-NGF in cell death (11), Nogo/Lingo-1/NgR in axonal growth (12, 13), and TrkA/NGF in survival and differentiation (14). p75 also undergoes shedding and receptor intramembrane proteolysis (RIP), resulting in the release of its intracellular domain (ICD), which itself possesses signaling capabilities (15–17).

Several lines of evidence implicate functional interactions between TrkA and p75NTR in NGF-triggered signal transduction (3, 18–20). TrkA and p75 receptors have nanomolar affinities for NGF and cooperate in transducing NGF signals (7, 21). The expression patterns of these two receptors overlap extensively (22) and in some instances, such as in the neurons of the dorsal root ganglion (DRG), TrkA is exclusively expressed in conjunction with p75 (23).

p75 has been experimentally demonstrated to enhance the response of TrkA to NGF (14, 24–26). In sympathetic neurons and oligodendrocytes, TrkA signaling inhibits the pro-apoptotic signaling of p75 (27–29). Primary DRG and sympathetic neurons derived from p75-null animals show attenuated survival responses to NGF (25, 26, 30), confirming the physiological role of p75/TrkA interactions. As the interaction between the two receptors seems to not engage the ligand binding domains of the extracellular region (31), the structural basis of such direct interaction is still unknown.

Here we demonstrate that the interaction between TrkA and p75 is mediated, at least in part, by the transmembrane domains. We validate these findings using functional studies in cells expressing the full-length receptors.

## RESULTS

### p75 and TrkA form a constitutive complex at the plasma membrane

We performed Förster Resonance Energy Transfer (FRET) experiments to determine if TrkA and p75 interact directly at the plasma membrane of live cells. HEK 293T cells transiently co-transfected with full-length TrkA tagged with mTurquoise (the donor fluorescent protein) and full-length p75 tagged with eYFP (the acceptor fluorescent protein) were imaged, and small regions of the plasma membrane were selected and analyzed. Illustrations of the TrkA-mTurq and p75-eYFP constructs used in FRET experiments are shown in Figure 1A. In each region of the cell membrane, we determined the FRET efficiency, the concentration of TrkA-mTurq and the concentration of p75-eYFP using the FSI-FRET software (Figure 1B) (32). These experiments were designed such that FRET can only occur between TrkA and p75, not between TrkA-TrkA or p75-p75. We also performed control FSI-FRET experiments using two unrelated proteins, LAT (Linker for the Activation of T-cells) and FGFR3 (Fibroblast Growth Factor Receptor 3), which are not expected to interact specifically and thus should give zero hetero-FRET. In addition, the proteins were designed such that the fluorescent tags are positioned differently with respect to the plasma membrane—the mTurq fluorophore is attached to the C-terminus of full-length LAT while the eYFP fluorophore is attached to the C-terminus of an FGFR3 construct lacking the intracellular region, “ECTM” (Figure 1A). Therefore, these two proteins will also not give rise to a non-specific FRET signal (random or “proximity” FRET) (33). As expected, due to the large separation between the fluorescent tags, the FRET efficiencies measured between these two control proteins are localized around zero at all concentrations measured (Figure 1B). Therefore, this control dataset demonstrates the scenario where there is no FRET between the proteins.

**Figure 1.**
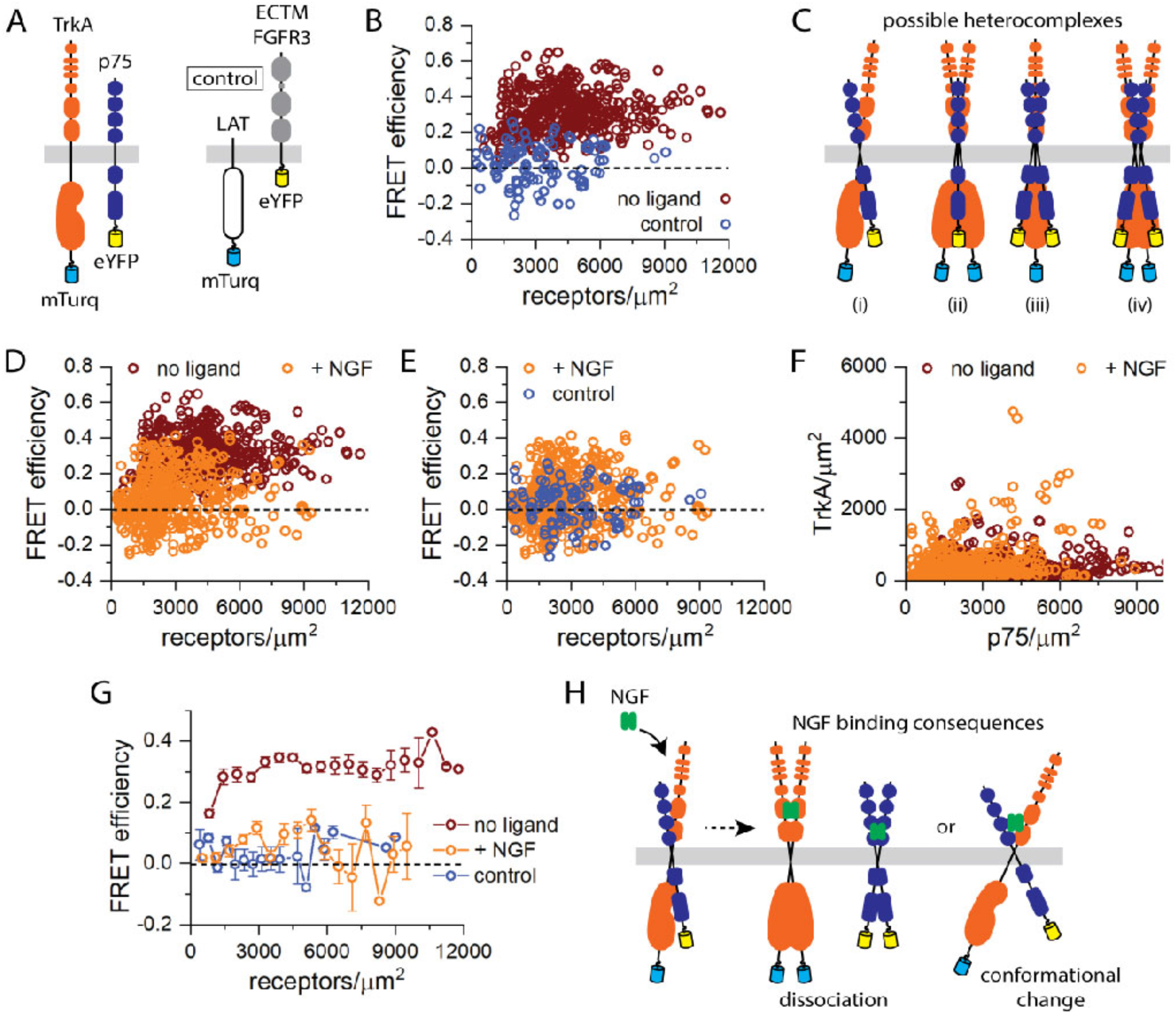
TrkA-p75 FSI-FRET experiments. (A) Illustrations of the TrkA-mTurq and p75-eYFP proteins used in FRET experiments along with the LAT-mTurq and ECTM FGFR3-eYFP proteins used in control experiments. (B) FRET efficiencies as a function of total receptor concentration measured for TrkA-mTurq and p75-eYFP in the absence of ligand compared to a zero FRET control dataset. (C) Illustrations of some possible stoichiometries of the TrkA-p75 heterocomplex: (i) heterodimer, (ii) heterotrimer of two TrkA and one p75, (iii) heterotrimer of one TrkA and two p75, (iv) heterotetramer or two TrkA and two p75. (D) FRET data for TrkA-mTurq and p75-eYFP in the presence of 100 ng/µL NGF compared to the data in the absence of NGF. (E) The FRET data for TrkA and p75 in the presence of NGF compared to the zero FRET control dataset. (F) Expression of TrkA-mTurq and p75-eYFP measured on the cell surface for the experiments performed in the absence and presence of NGF. (G) The FRET data for TrkA-p75 in the absence and presence of NGF and for the control dataset were binned and compared. (H) Illustrations of the possible consequences of NGF binding to the TrkA-p75 heterocomplex, which could be either dissociation of the heterocomplex to stabilize the respective homodimers or an NGF-induced conformational change.

In the absence of ligand, full-length TrkA and p75 exhibit positive (greater than zero) FRET efficiency values over all TrkA and p75 concentrations measured (Figure 1B). Therefore, this data suggests that TrkA and p75 interact directly at the plasma membrane. With this data alone, we cannot determine an accurate stoichiometry of the TrkA-p75 heterocomplex. Given that TrkA and p75 exist in monomer-dimer equilibrium in the absence of ligand, it is possible that TrkA and p75 associate as heterodimers, or oligomers of higher order (Figure 1C). Next, we sought to determine if NGF ligand binding influences the TrkA-p75 heterocomplex, and we performed similar FSI-FRET experiments for TrkA-p75 in the presence of 100 ng/µL NGF (Figure 1D). The FRET efficiencies measured for TrkA-p75 in the presence of NGF are noticeably lower compared to the data in the absence of ligand. Furthermore, comparison of the liganded TrkA-p75 FRET data to the LAT-FGFR3 control experiment data revealed no significant differences (Figure 1E), which suggests that the fluorophores attached to the C-termini of TrkA and p75 are too far away from one another to observe a FRET signal in the ligand-bound state. The expression levels of the TrkA and p75 at the cell surface are similar in both sets of experiments (+/- NGF) so these differences are not a reflection of altered gene expression (Figure 1F). The decrease in FRET may mean that the heterointeractions are abolished, for instance, due to ligand-induced homodimer stabilization, or it may be due to conformational changes in the heterocomplex which leads to decreased FRET.

The FRET data for TrkA-p75 in the absence and presence of NGF and the control dataset were binned and compared in order to visualize the average FRET efficiency as a function of receptor concentration (Figure 1G). For the control dataset and the TrkA-p75 data in the presence of NGF, the average FRET efficiencies remain around zero as expected from the raw data (Figure 1G). For the TrkA-p75 data in the absence of ligand, we observe average FRET efficiencies greater than zero over all concentrations (Figure 1G). Furthermore, at the low receptor concentration regime, the average FRET efficiencies increase as a function of receptor concentration, suggesting increasing TrkA-p75 interactions (Figure 1G).

### Direct interaction between p75 and TrkA transmembrane domains

Previous findings have suggested that TrkA can form a complex with p75-CTF (a membrane-anchored C-terminal fragment) by means of transmembrane domain interaction (17). In addition, the TM domain of p75 is involved in the formation of the high-affinity NGF binding sites (34), suggesting that the TM domain may mediate the direct interaction between p75 and TrkA. Therefore, we were interested in investigating the interaction between the p75 and TrkA, taking into account the recently reported NMR structures of p75 and TrkA TM domains (35, 36). We examined the interaction of p75-TM-wt with the TrkA-TM domain in lipid micelles using NMR spectroscopy. Increasing amounts of TrkA-TM were added to the ^15^N-labeled p75-TM in DPC micelles and the chemical shifts were monitored in a ^1^H-^15^N HSQC spectrum (Figure 2A). Chemical shifts are very sensitive to the electronic environment of a nucleus, and serve as an ideal instrument to probe the protein-protein interaction. Previous work in our laboratory found that p75-TM-wt forms spontaneous disulfide dimers (35). We titrated the ^15^N labeled p75-TM-wt disulfide dimer with increasing concentrations of TrkA-TM-wt solubilized in DPC micelles, retaining the constant lipid-to-protein ratio (LPR). The titration revealed no chemical shift changes. We used several LPRs and at least two independent preparations of p75-TM-wt. As the dimerization of p75-TMD-wt is irreversible (35) we performed the experiments with the mutant p75-C257A, which forms non-covalent homodimers (35) and allows the possibility to obtain the monomeric p75-TMD. According to the previous work (35), the C257A mutation does not induce any substantial changes to the structure of p75 TMD. Several chemical shift changes were observed in the HSQC-NMR spectrum of p75-TM-C257A upon titration with TrkA-TM-wt, suggesting the formation of specific p75/TrkA heterocomplexes (Figure 2B).

**Figure 2.**
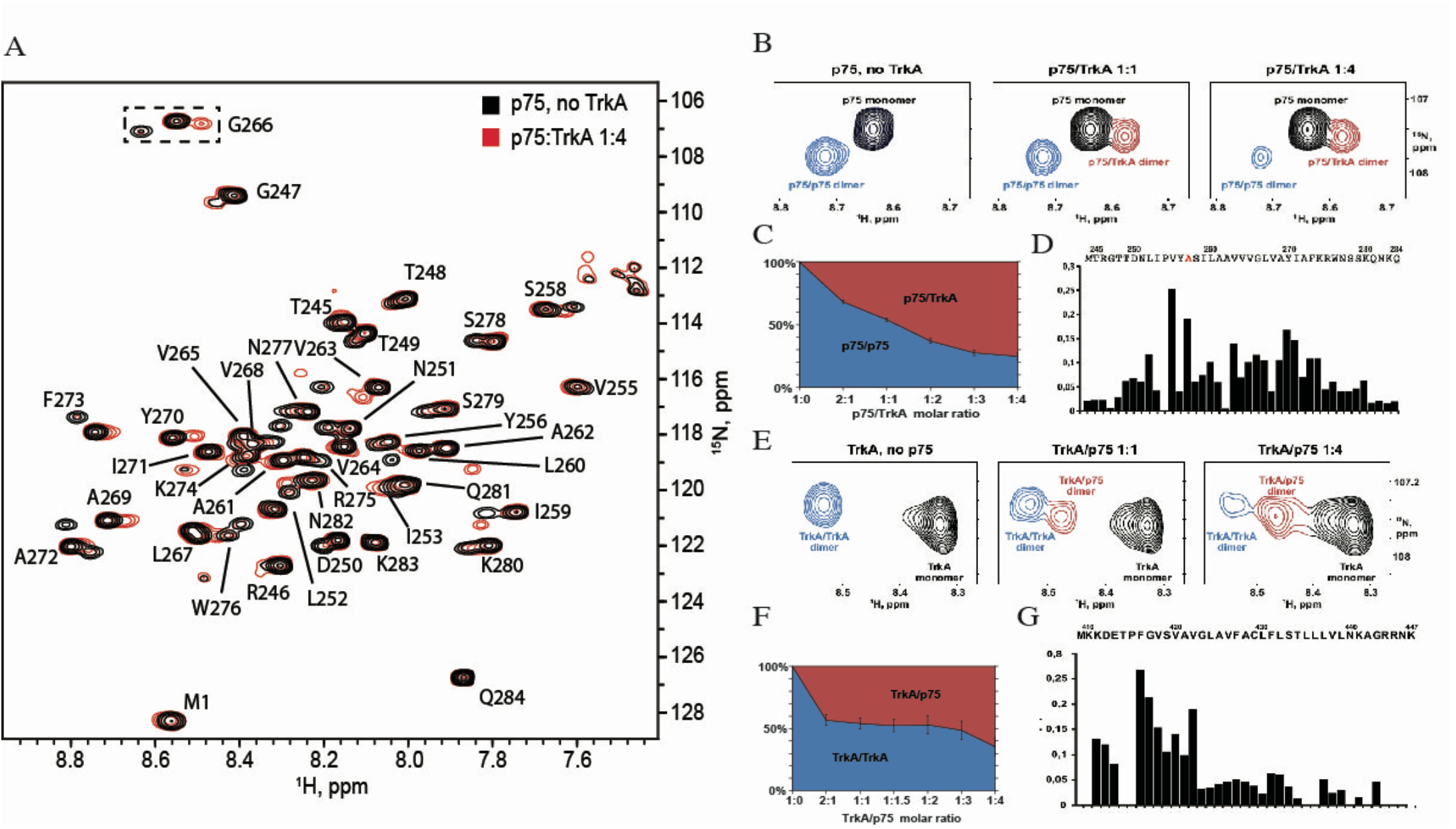
p75/TrkA interactions as observed by NMR. A) Overlay of two ^15^N-TROSY experiments: (black) ^15^N-labeled p75 without TrkA and (red) ^15^N-labeled p75 after addition of unlabeled TrkA with p75:TrkA molar ratio 1:4. ^1^H-^15^N assignments of p75 backbone amid proton resonances are provided. B) ^15^N-labeled p75-TM-C257A titration with unlabeled TrkA TM. Left to right: p75 monomer (black), p75-p75 homodimer (blue) and p75-TrkA heterodimer (red) states are observed in the G266 amide proton cross-peak in ^1^H/^15^N-HSQC spectra. G266 was chosen as representative as its cross-peak is situated away from other peaks and it shows clear monomer-homodimer-heterodimer transitions. C) Chemical shift changes observed upon interaction with TrkA are shown on top of p75-TM sequence. D) Population of p75-p75 homodimers relative to that of p75-TrkA heterodimers (p75-p75 peak intensity is divided by sum of p75-p75 and p75-TrkA peak intensities), expressed as a function of the p75/TrkA molar ratio. The population of p75-p75 dimer decreases while that of p75-TrkA dimer increases as more TrkA is added to the sample. For all experiments the lipid to protein molar ratio (LPR) remains constant at 80. E) ^15^N-labeled TrkA-TMD titration with unlabeled p75-TM-C257A. Left to right: TrkA monomer (black), TrkA-TrkA homodimer (blue) and p75-TrkA heterodimer (red) states are observed in the amide proton cross-peak in ^1^H/^15^N-HSQC spectra. F) Chemical shift changes observed upon interaction with p75 are shown on top of TrkA-TMD sequence. G) Population of TrkA-TrkA homodimers relative to that of TrkA-p75 heterodimers (TrkA-TrkA peak intensity is divided by sum of TrkA-TrkA and TrkA-p75 peak intensities), expressed as a function of the TrkA/p75 molar ratio. The population of TrkA-TrkA dimer decreases while that of TrkA-p75 dimer increases as more p75 is added to the sample.

To identify the oligomer size of the complex, we measured the cross-correlated relaxation rates of p75-TM-C257A signals (Figure S1). According to the recent work, the NMR-derived hydrodynamic radii of TM domains in DPC micelles can be used to distinguish the various oligomeric forms of the proteins (37). Here we observed the rotational correlation time (and hydrodynamic radius) of a p75-TM-C257A monomer at 45 °C to be 10.2±0.4 ns (2.61 nm), a p75-TM-C257A homodimer to be 13.1±0.6 ns (2.85 nm) and the heterocomplex to be 12.7±0.8 ns (2.82 nm). In other words, the observed new complex formed by TrkA-TM and p75-TM-C257A is a heterodimer as the rotational correlation time of the heterocomplex was similar to that of the homodimer.

With the increase of TrkA concentration, the percentage of p75 homodimer decreased while that of p75/TrkA heterodimer increased (Figure 2C). This implies that homo- and heterodimerization of p75-TM are the competing processes. The titration curve revealed homodimerization and heterodimerization constants of comparable magnitudes. Similar effects were observed when ^15^N-labeled TrkA-TMD sample was titrated with the unlabeled p75-TM-C257A (Figure 2E, F). Addition of p75-TM-C257A decreased the concentration of TrkA-TM homodimer, while the novel heterodimeric state had emerged, which is indicative of the competition. Thus, we can state that TrkA interacts with the monomeric form of p75 TM domain but does not bind the disulfide-crosslinked dimer of the protein. Most likely, the covalent dimerization shields some of the p75 residues necessary to interact with the TrkA TM domain, or the interaction requires a rearrangement of the dimer that cannot be achieved due to the restraints imposed by the disulfide bonds.

Chemical shift (CS) changes were detected along the p75 TMD sequence (Figure 2D), which is expected as the TrkA interaction breaks the p75-TM-C257A dimerization. The residues with the highest chemical shift changes are shown in the Figure 2D. To find the residues undergoing chemical shifts changes in the TrkA-TMD, we performed the titration on labeled ^15^N-TrkA-TMD homodimer with unlabeled p75-TM-C257A (Figures 2E). With increasing p75-TM-C257A concentration, the percentage of TrkA homodimer decreased while the heterodimer increased (Figure 2F). The NMR chemical shifts indicated that the region of higher CS changes (Figure 2G, Δδ>0.1) upon interaction of p75-TMD is located mainly at the N-terminus of TrkA TMD.

These results support a direct interaction between p75-TMD and TrkA-TMD and suggest that the formation of a heterodimer outcompetes the homodimerization of each TM domain. Although the NMR shows that the interaction is direct, we cannot use the CS changes to identify the protein-protein interface between the TM domains in a membrane. Recently it has been shown that, by contrast to soluble proteins, CS changes have almost zero predictive power to map protein interfaces in transmembrane regions (38). CS changes primarily report hydrogen bonding and are insensitive to van-der-Waals contacts between the protein side chains, which are the main driving force for dimerization of membrane proteins (38).

### Multiscale Molecular Dynamics

The crowding of the NMR spectra with several TrkA and p75 species (monomer, homo- and heterodimers) precludes the complete CS assignment and the structure calculation of the heterocomplex. To explore further the interaction between TrkA and p75 TMDs we used molecular dynamics (MD) (Table 1 and Figure 3). MD simulations provide a useful approach for modeling the transmembrane domain interactions (39). Both full-atom (FA) and coarse-grained (CG) modeling has been previously used to optimize the dynamics and interactions between different transmembrane domains (39). To model the heterodimerization of p75-TMD and TrkA-TMD, two CG helices were inserted in a parallel orientation relative to one another separated 6 nm in a preformed POPC bilayer and 24 simulations of 5 µs were run (total time 120 µs) (Table S1 and Figure 3A). In all but one of the 12 simulations, the TrkA/p75 heterodimer was formed within the first 2 µs (except for the simulation number #5 that formed the heterodimer at 5 µs) and did not dissociate during the remainder of the simulation (Figure 3B). The POPC model membrane was well equilibrated with average values for the area per lipid and hydrophobic thickness (between glycerol groups) of 63.2 Å2 and 34.8 Å respectively that are in good agreement with the experimental values (40) (Table S2). From each of the heterocomplexes (Figure 3C), we compute the root mean squared deviation (RMSD) between them and found a cluster of 7 models with an average RMSD of 2.12 Å (Figure 3D). The final model was converted from coarse-grained to full-atom to further study the packing of the interaction in a POPC lipid bilayer during 100 ns of FA-MD, done in triplicate. The final POPC model membrane was well equilibrated with average values for the area per lipid and hydrophobic thickness (between phosphate groups) of 63.3 Å2 and 38.4 Å respectively that are in good agreement with the experimental values (40). The membrane electron density was calculated and shown in the Figure S2. The interhelix distance between residues at the C-terminus of the helix (p75-W276 and TrkA-K441) was calculated along the total simulation time (Figure 3E), indicating the equilibration of a stable complex.

**Figure 3.**
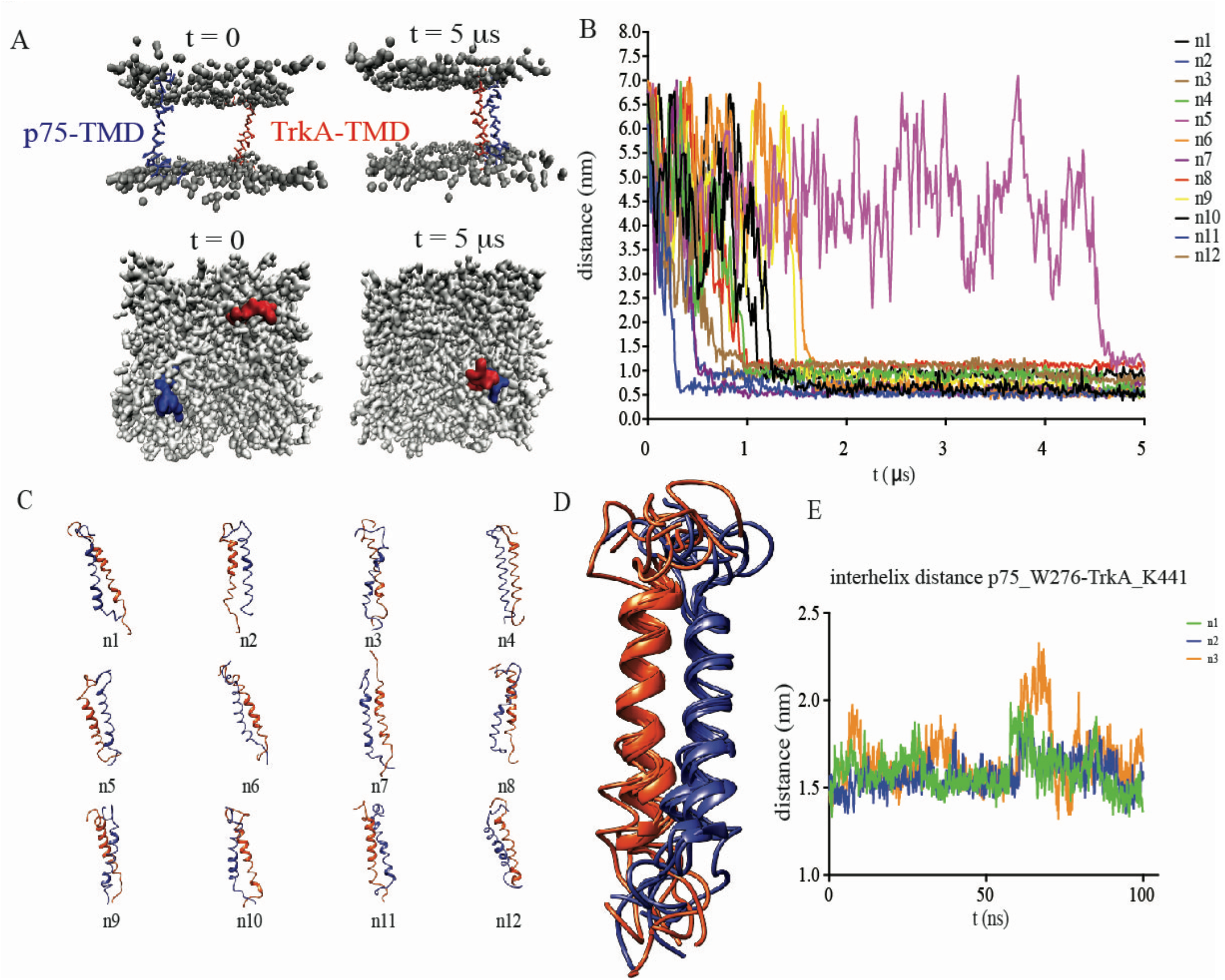
Multiscale Molecular dynamics of TrkA-TMD and p75-TMD. A) Coarse-grained TrkA-TMD and p75-TMD helix dimerization simulation. The initial system configuration (0 µs) consists of two helices (red and blue) inserted in a POPC bilayer in a parallel orientation with an interhelix separation of dHH ≈ 55 Å. The choline, phosphate and glycerol (gray) backbone particles of the POPC molecules are shown. The snapshot at 5 µs illustrates the stable TM helix heterodimer. B) Distance between TrkA-TMD and p75-TMD during CG-MD simulation time. C) Structural models of the final conformations from the 12 simulations. In blue p75 and in red TrkA is shown. D) Superposition of the 7 conformations with lowest rmsd found by CG-MD. E) Interhelical distance between p75-TMD-W276 and TrkA-TMD-K441 in the FA-MD simulation done by triplicate.

The protein interface of p75-TMD participating in the interaction with TrkA-TMD is C_257_S_258_xxA_261_A_262_xxV_265_G_266_xxA_269_xx (Figures 4A and 4B). This interface contains the motif A_262_xxxG_266_xxA_269_ that was previously identified in the homodimerization of p75-TMD-C257A (35) and is supported by the NMR experiments shown above, indicating that heterodimerization with TrkA-TMD competes with the p75-TMD non-covalent homodimerization. In addition, the residue C257 forms a part of the heterodimer interface supporting our observations that disulfide dimers do not significantly bind to the TrkA-TMD. The TrkA-TMD heterodimer interface is formed by the motif V_418_xxxV_422_xxxV_426_F_427_xxL_430_ (Figure 4B) where the central valine residues make the closest contact with the p75-TMD. Interestingly, several of these residues are conserved in TrkB and TrkC (Figure 4C) suggesting that these receptors interact with p75 in a similar manner as TrkA.

**Figure 4.**
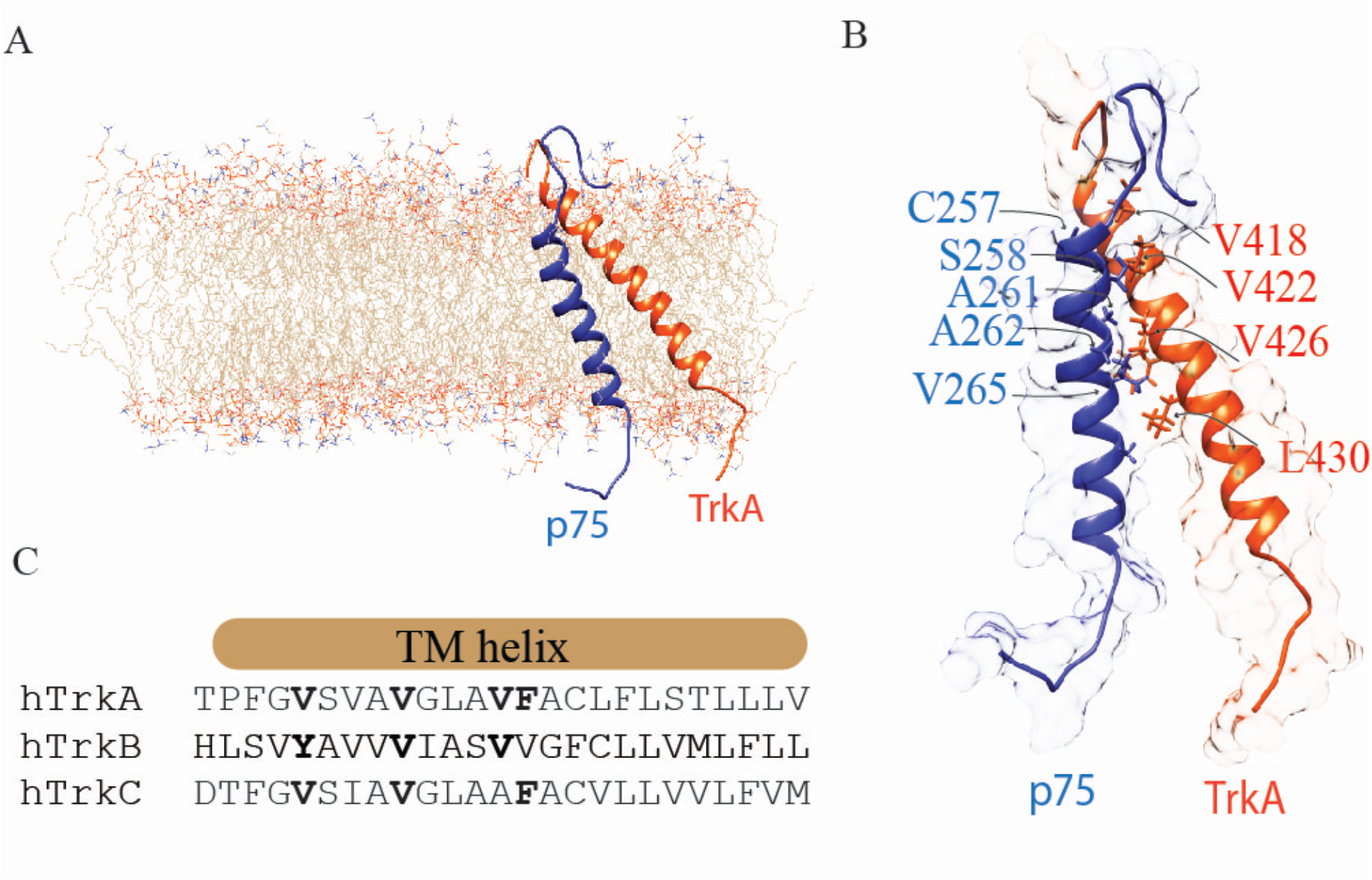
Structural models of the p75/TrkA TMD heterodimer. A-B) Schematic representation of the spatial structure of the heterodimer p75-TMD (blue) and TrkA-TMD (orange) after 100 ns full-atom MD. The residues participating in the dimer interface are shown by blue (p75) and red (TrkA). C) Protein sequence alignement of TrkA, TrkB and TrkC TMDs. In bold the conserved residues.

Altogether, the NMR and FRET data support the direct interaction between TrkA and p75 and the MD provides insight into a possible heterodimer model.

### The transmembrane heterodimer interface modulates TrkA activation and sensitization to lower concentrations of NGF

*In vivo* data suggest that in sensory neurons p75 helps TrkA to respond to the lower concentrations of NGF (26) and enhances the response of TrkA to NGF (14, 24). One current hypothesis is that the binding of p75 to TrkA induces a conformational change in TrkA that facilitates both the binding of NGF to TrkA (24) and the activation of TrkA (26). To test if the protein interface found above has any physiological role we sought to determine if mutations on the p75 transmembrane protein interface influences TrkA activation to lower concentrations of NGF (Figure 5A). We co-expressed p75 with TrkA full-length receptors in Hela cells and stimulated with increasing concentrations of NGF (0, 0.1, 1, 10 and 100 ng/mL). Western blot of cell lysates were probed with specific antibodies against the activation loop of the TrkA kinase domain (Tyr675 and Tyr676) (Figure 5B). Quantification of the protein bands corresponding to the phosphorylation of TrkA was plotted against NGF concentration. Fitting the data to a dose (NGF)–response (phosphorylation) curve allows an estimation of the EC50 of NGF, the concentration of NGF that provokes a response half way between the basal response and the maximal response (Figure 5D). Hela cells transfected with TrkA present a LogEC_50_ of −9.219 ± 0.087 (an EC_50_ = 6.03 x10-^10^ M). In cells co-expressing TrkA and p75 an LogEC_50_ of −9.524 ± 0.176 (an EC_50_ = 2.99×10^−10^ M) was found, showing a small, but significant effect of p75 on the activation of TrkA by NGF. The parallel curve suggested an agonist effect of p75 and NGF on the activation of TrkA. To analyze the effect of p75-TMD we used a construct of p75 with its transmembrane domain swapped with the one from the tumor necrosis factor receptor (TNFR), mutant p75-TNFR. A decrease in the NGF sensitivity was observed in comparison to p75-wt (LogEC50 −8.56 ± 0.54, EC_50_= 2.7×10^−9^ M), indicating that the effect of p75-wt is lost in the p75-TNFR construct. As the protein heterodimer interface contains the motif A_261_A_262_xxxG_266_xxA_269_ we made a construct with a triple mutation A262,G266,A269 to Ile (p75-AGA mutation). The rationale behind this is that the introduction of a hydrophobic bulky residue, Ile, would impair the proper interaction with the TrkA-TMD. Fitting of the values obtained from the lysates transfected with TrkA and p75-AGA showed a LogEC50 of −8.776 ± 0.037, that corresponds to an EC50 = 1.7×10^−9^ M (Figure 5D), that accounts for more than one order of magnitude higher than in the presence of p75-wt supporting that this interface plays a key role in TrkA activity modulation by p75.

**Figure 5.**
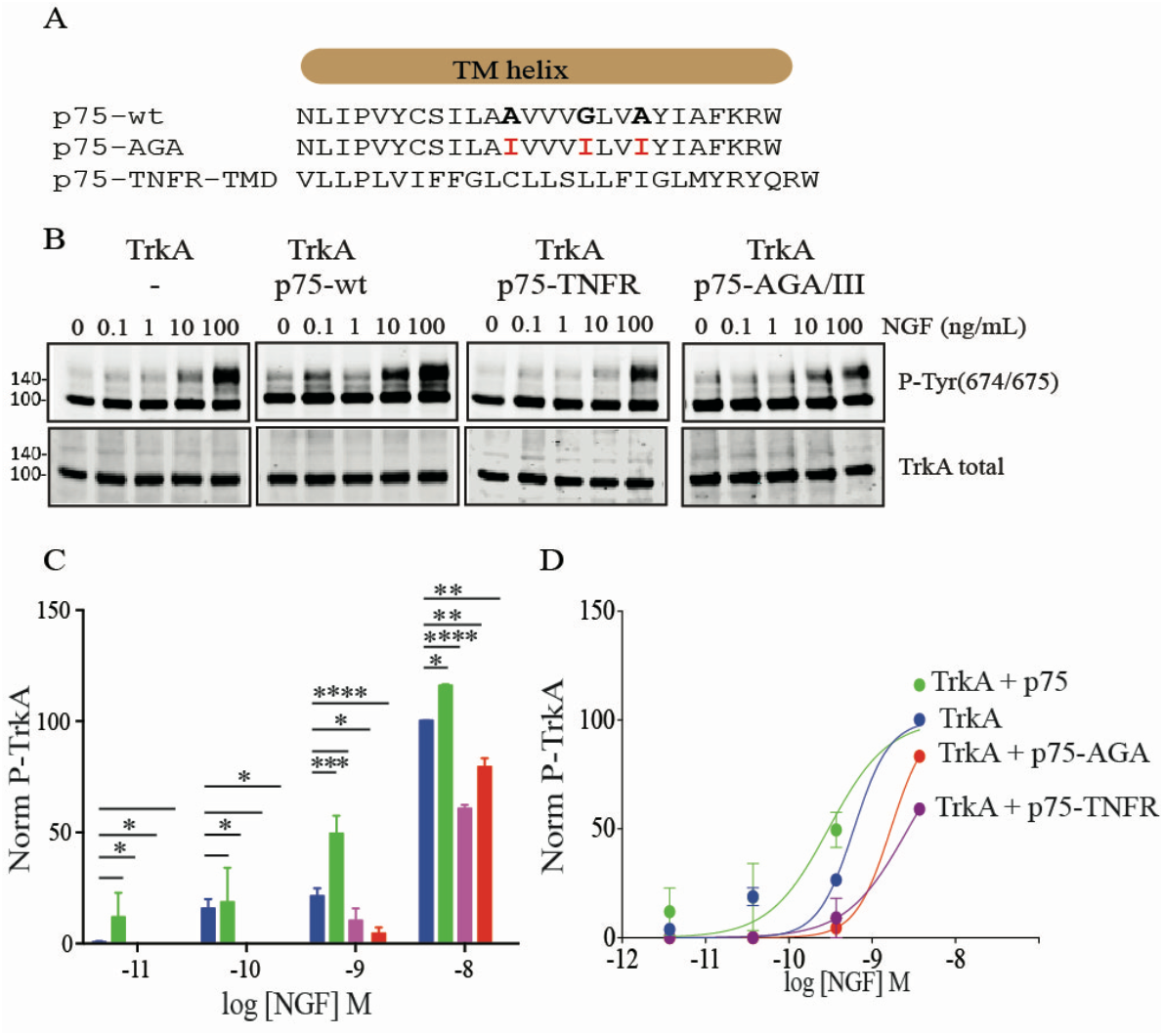
TrkA activation is modulated by p75-TMD. A) Protein sequences alignment of the different mutant constructs of p75-TMD. The residue mutated is shown in bold. B) Western blots of lysates from Hela cells transfected with the indicated constructs and stimulated with increasing concentrations of NGF. Membranes were probed using a TrkA-P-Tyr675 specific antibody. C-D) Normalized activation of TrkA using increasing concentrations of NGF in the absence of the presence of p75 mutant constructs indicated. Bars represent the standard error of at least three independent experiments. P values are reported in the text.

### p75 needs a specific interface in the transmembrane domain to interact to TrkA

The finding that the activation of TrkA in the presence of the p75-AGA mutant is lower than in the absence of p75 suggested an antagonist or inhibitor behavior for this mutant. To further study the effect of this mutation on the heterodimer complex, we introduced the triple mutation AGA/III into the p75-TMD and performed a CG-MD followed by FA-MD simulation similar to the p75-TMD-wt constructs shown above (Figure S3). MD analysis showed that although p75-TMD-AGA mutant still interacts and binds to the TrkA-TMD with similar kinetics as the p75-TMD-wt, the heterodimer arrangement is changed significantly. It has been previously shown that TrkA-TMD contains two homodimer interfaces; an active dimer formed upon NGF binding and an inactive dimer formed in the absence of NGF. The 12 independent simulations of p75-TMD-wt showed a restricted binding interface localized close to the inactive homodimer interface, leaving the active homodimer interface of TrkA free and accessible (Figure 6A). However, after 12 independent simulations the end-point of p75-TMD-AGA is almost equally distributed in all the possible TrkA-TMD interfaces (Figure 6B), where the active homodimer interface is hidden by p75-TMD. This result indicates that p75-TMD-AGA could impair TrkA active homodimerization and may explain the weaker activation of TrkA in the presence of p75-TMD-AGA.

**Figure 6.**
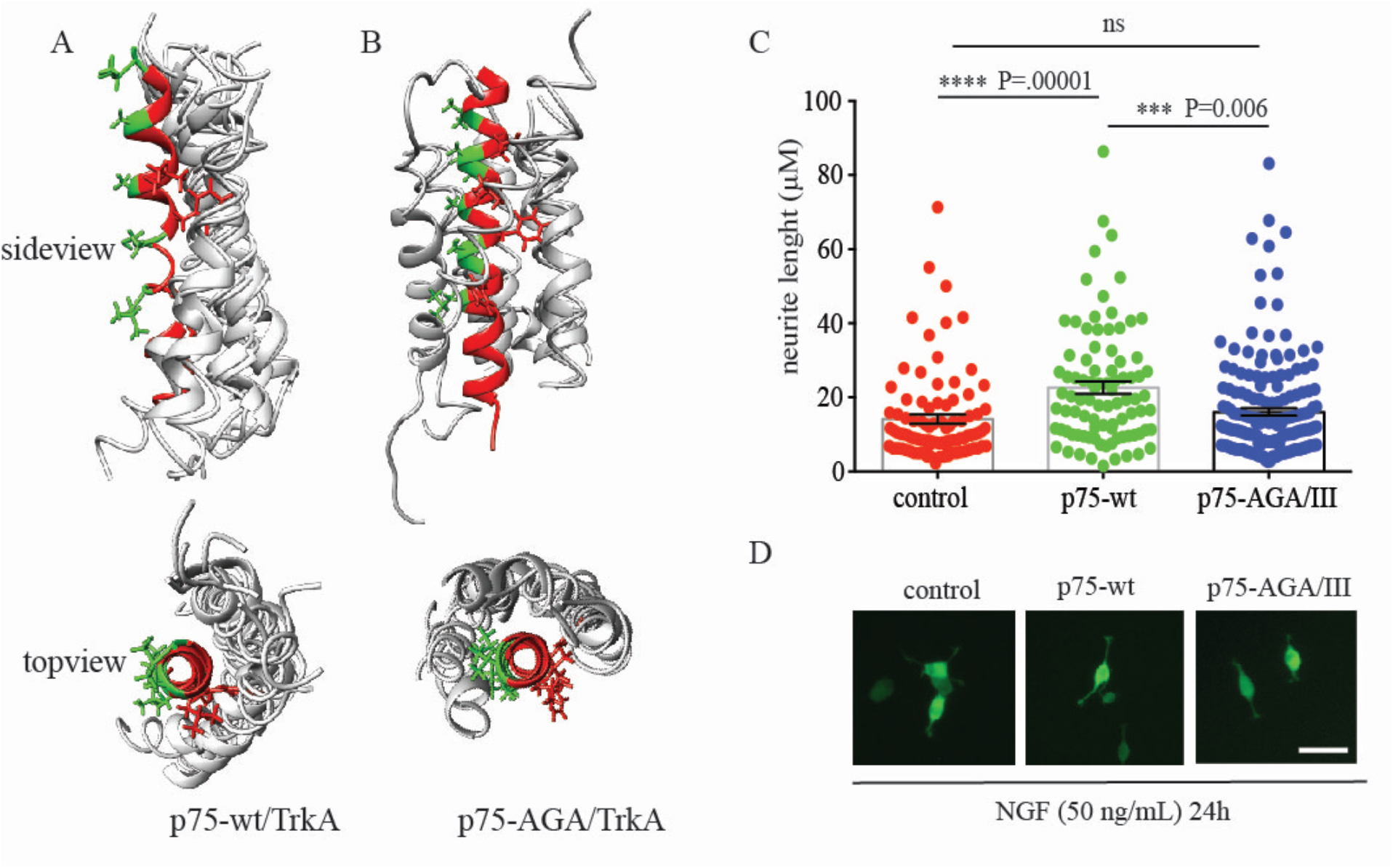
Effect of the mutation of the p75 heterodimer interface. A-B) Result of 12 simulations by CG-MD of p75-TMD-AGA mutant (A) or p75-TMD-wt (B) and TrkA-TMD in POPC model membranes. The position of the p75-TMD helix (gray) respect to the TrkA-TMD (red) after each simulation is shown. In green and red are shown the residues that belong to the active and inactive homodimer interface of TrkA described in Franco et al. C) Quantification of the neurite length (µm) of PC12 cells electroporated with the indicated constructs and GFP at 24h of addition of NGF (50 ng/mL). Bars represent the standard error of at least three independent electroporation experiments. Statistical analysis was performed with one-way Anova analysis was used and the P values are reported above each bar. B) Representative fluorescence microscopy of PC12 cells electroporated with the indicated constructs stimulated with NGF (50 ng/mL) for 24 hours post-electroporation. Bar represents 50 µm.

### p75-AGA/III reduces NGF-induced differentiation of PC12 cells

To further support our finding that p75 needs a specific heterodimer interface to fully activate TrkA, we overexpressed p75-wt and p75-AGA/III in PC12 cells that endogenously express TrkA and quantified the neurite length upon stimulation with NGF. As shown in Figure 7A, the PC12 cells transfected with p75-AGA/III had shorter neurite lengths at 24h than cells transfected with p75-wt (16.02 µm ± 0.98, n=181 vs 22.63 µm ± 1.69, n=89) and similar length as PC12 cells transfected with the empty vector (14.09 µm ± 1.25, n=92) (control in Figure 7A). These experiments suggest a reduction in the activation of TrkA by NGF of p75-AGA/III in comparison to p75-wt.

## Discussion

The present study provides, to the best of our knowledge, the first structural evidence of a direct interaction between p75 and TrkA. While data from *in vitro* and *in vivo* experiments has suggested the existence of a complex formed by p75 and TrkA (41–43), repeated attempts to observe the direct interaction between both receptors using different biochemical and structural approaches have been unsuccessful. Experimental evidence of the existence of a TrkA/p75 complex were based on co-immunoprecipitation studies (17, 21, 44) and by biophysical methods such as co-patching (45) and fluorescence recovery after photobleaching (46). In addition, a handful of studies have suggested that the transmembrane and intracellular domains of p75 could be responsible for its interaction with TrkA (21, 34, 47, 48).

Here we demonstrated that the complex formed by p75 and TrkA is mediated by the TM domains, supporting the findings by previous reports (21, 34). The results of our NMR titration experiments point to a relatively weak affinity constant, similar to that calculated for p75 non-covalent dimerization. This is around 10 times weaker than the affinity constant calculated for glychopohrin-A homodimerization, and explains why these complexes have been difficult to detect by co-immunoprecipitation in the presence of detergents (i.e, glycophorin A TM domain dimers are resistant to SDS-PAGE). Hetero-crosslinking experiments similarly failed to detect p75/TrkA complexes, although probably for different reasons, as crosslinking requires the specific residues (i.e Lys) to be close to each other and oriented in a specific manner, not always possible even in a heterocomplex. Our results are in agreement with those of fluorescence recovery after photobleaching (FRAP) experiments, which show that p75 is fully mobile at the cell membrane but becomes restricted in mobility upon TrkA co-expression (46), and with biochemical evidence suggesting that the TM domain of p75 is necessary for the formation of high-affinity NGF binding sites (34). Although the TMD interaction is weak, *in vivo* the levels of p75 and Trk normally exist at a ratio of approximately 10:l (49, 50) favoring their heterointeractions interaction over TrkA homointeractions.

Recently it has been shown that TrkA has two homodimer interfaces in the TMD; one active and one inactive (36). The active interface corresponds to the TrkA bound to its ligand NGF. And the inactive dimer interface corresponds to the pre-formed dimer of TrkA in the absence of NGF. The observed binding of p75-TMD to TrkA takes place mainly through an interface that is opposite to the active interface and partially covering part of the inactive dimer interface, suggesting that binding of p75 to TrkA may favor the formation or stabilization of TrkA active homodimers. In addition, stabilization of a pre-formed dimer would be compatible to an increase in the affinity of TrkA for NGF in the presence of p75 (18), suggesting that the heterodimer p75/TrkA described here forms the basic unit of the NGF high-affinity sites. The finding that mutations in the p75 protein interface, as shown here with the p75-AGA mutant, impact the TrkA activation and supports the requirement of specific TMD interactions in the neurotrophin receptors. As it has been shown recently, NGF binding can induce the rotation of the TrkA TM dimer form the inactive to the active interface (36, 51). This conformational change is supported by our FRET analysis, which reveals that NGF binding alters the TrkA-p75 heterocomplex that we observed in the absence of ligand. There are some possible explanations for this result, which are both illustrated in Figure 1H. The first option is that NGF binding could cause the dissociation of the TrkA-p75 heterocomplex, stabilizing the respective homodimers instead. Another explanation is that NGF binding induces a conformational change of the TrkA-p75 heterocomplex that alters the positioning of the fluorescent proteins, increasing their separation and thus decreasing the FRET signal. While this data cannot distinguish between these two possible effects, the FRET data clearly demonstrate that TrkA and p75 interact directly in the absence of ligand and that NGF binding alters the heterocomplex.

Our MD analysis of p75-AGA/TrkA interactions showed that the inactive dimer interface is accessible suggesting that p75-AGA interaction may displace the equilibrium towards the inactive homodimer of TrkA in the absence of NGF. This would affect the activation of TrkA and lead to lower cell differentiation capabilities of PC12 cells overexpressing the p75-AGA mutant. Alternatively, the binding of the p75-AGA mutant may affect the conformational change induced by NGF binding resulting in a less activation of TrkA.

Altogether, we show that a specific transmembrane interaction is required for the positive role of p75 in TrkA activation by NGF. In conclusion, we provide a new structural insight on the highly dynamic p75/TrkA heterocomplex, paving the way to new investigations about the biological relevance of such interactions.

## EXPERIMENTAL PROCEDURES

### p75-TM and TrkA-TM constructs for cell-free expression

The gene encoding transmembrane and juxtamembrane residues 245-284 (MT245RGTTDNLIPVYCSILAAVVVGLVAYIAFKRWNSSKQNKQ284) of human p75 receptor (p75-TM-wt) was amplified by PCR from six chemically synthesized oligonucleotides (Evrogen, Russia) partially overlapped along its sequence. The C257A point mutant form of p75TM (p75-TM-C257A) was obtained by site-directed mutagenesis by PCR. The PCR products were cloned into a pGEMEX-1 vector by three-component ligation using the NdeI, AatII and BamHI restriction sites. Expression constructs for human TrkA-TM (MK410KDETPFGVSVAVGLAVFACLFLSTLLLVLNK<u>A</u>GRRNK447) were similarly prepared by PCR.

### Fully Quantified Spectral Imaging (FSI)-FRET experiments

Human embryonic kidney (HEK) 293T cells used in the FRET experiments were purchased from American Type Culture Collection (Manassas, VA; CRL-3216). The cells were cultured at 37 °C and 5% CO2 in Dulbecco’s Modified Eagle Medium (DMEM; Thermo Scientific; 31600-034) containing 3.5 g/L D-glucose, 1.5 g/L sodium bicarbonate, and 10% fetal bovine serum (FBS; Sigma-Aldrich; F4135). HEK293T cells were seeded in 35 mm glass bottom collagen-coated petri dishes (MatTek Corporation, MA) at a density of 2 × 10^5^ cells/dish and cultured for 24 hours. The cells were co-transfected with pcDNA constructs encoding for TrkA tagged with mTurquoise (mTurq, the donor) and p75 tagged with enhanced yellow fluorescent protein (eYFP, the acceptor). The TrkA-mTurq plasmid was generated as described (32, 52). The p75-eYFP construct was cloned by overlapping PCR into the same pcDNA vector. The LAT and ECTM FGFR3 plasmids used for control experiments were generated as described previously (53, 54). Transfection was performed with Lipofectamine 3000 (Invitrogen, CA; L3000008) using 1-4 µg of total DNA at a TrkA:p75 ratio of 2:1 or 1:1. In addition, cells singly transfected with either TrkA-mTurq or p75-eYFP were used for calibration as described (32). After twelve hours following transfection, the cells were washed twice with starvation media (serum-free, phenol red-free media) and serum-starved in starvation media for 12 hours overnight. Prior to imaging, the starvation media was replaced with hypo-osmotic media (10% starvation media, 90% diH2O, 25 mM HEPES) to ‘unwrinkle’ the highly ruffled cell membrane under reversible conditions as described (55). Cells were incubated for 10 minutes and then imaged under these conditions for approximately 1 hour. In some experiments, soluble human beta nerve growth factor (hβ-NGF; Cell Signaling Technology; 5221SC) was diluted to a final concentration of 100 ng/µl with the hypo-osmotic media before adding to the cells.

Cell images were obtained following published protocols (32) with a spectrally resolved two-photon microscope set up using a Zeiss Inverted Axio Observer and the OptiMis True Line Spectral Imaging system (Aurora Spectral Technologies, WI) with line-scanning capabilities (56, 57). Fluorophores were excited with a mode-locked laser (MaiTai™, Spectra-Physics, Santa Clara, CA) that generates femtosecond pulses between wavelengths 690 nm to 1040 nm. For each cell, two images were collected: the first at 840 nm to excite the donor and the second at 960 nm to primarily excite the acceptor. Solutions of purified soluble fluorescent proteins (mTurq and eYFP) were produced at known concentrations following a published protocol (58) and imaged at each of these excitation wavelengths. A linear fit generated from the pixel-level intensities of the solution standards was used to calibrate the effective three-dimensional protein concentration which can be converted into two-dimensional membrane protein concentrations in the cell membrane as described (32). Small micron sized regions of the cell membrane were selected and the FRET efficiency, the concentration of TrkA-mTurq, and the concentration of p75-eYFP present in the cell membrane were quantified using the FSI-FRET software (32).

### Cell-free gene expression

Bacterial S30 cell-free extract was prepared from 10 L of cell culture of the *E. coli* Rosetta(DE3)pLysS strain, using a previously described protocol (Aoki et al., 2009; Kai et al., 2012; Schwarz et al., 2007). Preparative-scale reactions (2-3 mL of reaction mixture) were carried out in 50-mL tubes.

### Titration of TrkA and p75 transmembrane domains by NMR

All TrkA/p75 titration 15N-TROSY experiments were carried out at LPR 80, pH 5.9, temperature 318K with 20 mM NaPi buffer. Two independent sets of experiments were conducted: (1) unlabeled p75-TM-C257A was incrementally added to 0.5 mM sample of 15N-labeled TrkA-TM, and (2) unlabeled TrkA-TM was incrementally added to the 0.4 mM sample of 15N-labeled p75-C257A-TM sample to observe p75-TrkA interactions. Intensities of corresponding peaks were measured at each point, population of the p75-p75 dimer, TrkA-TrkA dimer and TrkA/p75 complex were calculated and plotted against TrkA/p75 molar ratio.

### Modulation of TrkA activity by p75

Hela cells were transfected with 1µg of TrkA and 1µg of p75 or p75-TNFR using PEI (ratio 10:1). 24 hours after transfections cells were lifted and split in identical numbers to a 6 well plate. 48 hours after transfection were starved for 2 hours with DMEM without serum and stimulated with different concentrations of NGF (from 0 to 100 ng/mL) for 15 minutes. Cells were washed with PBS and lysed with TNE buffer on ice for 15 minutes. Lysates were clarified by centrifugation and the cell supernatants quantified and analyzed by SDS-PAGE western immunoblots. Phospho-Tyrosine specific antibodies (anti P-Tyr674/675 from Cell Signalling 1:3000) and anti-p75 intracellular antibody (Promega) were used. To quantify the effect of p75 on TrkA we consider an allosteric interaction between p75 and TrkA and fit to a dose/response curve. The protein band corresponding to the phospho-Tyr signal was quantified and the ratio to the total TrkA was calculated. This is the response in the Figure 4. We plot the log of the concentration of NGF versus the response and the curve was fit to a log(agonist) vs response (three parameters) equation using the GraphPad software. The equation is Y=Bottom + (Top-Bottom)/(1+10^((LogEC50-X))), and the EC50 is the concentration of agonist, in this case NGF, that gives a response half way between Bottom and Top. At least three independent experiments were quantified.

### Coarse Grained Molecular Simulation Methods

One monomer from the TrkA-TMD dimer structure (PDB:2n90) and one monomer from the p75-TMD dimer structure (PDB:2mic) were converted to a coarse grained CG model using the script martinize.py from the martini web page (www.cgmartini.nl) and the tools from Gromacs 5.0.5. In CG models 4 heavy atoms are grouped together in one coarse-grain bead. Each residue has one backbone bead and zero to four side-chain beads depending on the residue type (Monticelli, Kandasamy et al., 2008). For all helix dimerization simulations, two α-helices were inserted into a preformed 1-palmitoyl-2-oleoyl-sn-glycero-3-phosphocholine (POPC) bilayer (containing 260 lipids) such that they were separated by an interhelix distance dHH ≈ 55 Å (Figure 6). Each system was solvated with 2975 CG water particles and 0.15 M NaCl counter ions. The energy of the system was minimized and followed by 12 MD simulations of 5 µs each simulation, total time 60 µs. CG simulations were performed using GROMACS v 5.0.5 (www.gromacs.org) (Van Der Spoel, Lindahl et al., 2005). All simulations were performed at constant temperature, pressure, and number of particles. The temperatures of the protein, lipid, and solvent were each coupled separately using the Berendsen algorithm at 305 K, with 774 a coupling constant τT = 1 ps. The system pressure was semiisotropically using the Parrinello-Rahman algorithm at 1 bar with a coupling constant τP = 12 ps and a compressibility of 3 Å~ 10−4 bar−1. The time step for integration was 20 fs. Coordinates were saved for subsequent analysis every 200 ps.

### Atomic Molecular Dynamics

GROMACS 5.0.5 was also used for all full atom MD simulations. CG models were converted to FA using the CHARMM-GUI portal (www.charmm-gui.org). FA was calculated using the CHARMM36m force field. Long-range electrostatics was calculated using the particle mesh Ewald method with a real-space cutoff of 10 Å. For the van der Waals interactions, a cutoff of 10 Å was used. The simulations were performed at a temperature of 303.15 K using a Nose-Hoover thermostat with τT = 1 ps. A constant pressure of 1 bar was maintained with a Parrinello-Rahman algorithm with an semiisotropic coupling constant τP = 5.0 ps and compressibility = 4.5 Å~ 10−5 bar−1. The integration time step was 2 fs. The LINCS method was used to constrain bond lengths. Coordinates were saved every 5 ps for analysis. Analysis of all simulations was performed using the GROMACS suite of programs. VMD (Humphrey, Dalke et al.,1996) and Chimera UCSF (Pettersen, Goddard et al., 2004) were used for visualization and graphics. Membrane equilibration was assed measuring the area per lipid and the membrane thickness using the APLVoro application (59). The electron density profiles were calculated using the *gmx density* tool in Gromacs. A representation of the electron density of the POC model membrane with TrkA and p75 TMDs is shown in the Figure S2.

### Cell culture and transfection

Hela cells, which do not endogenously express neither TrkA nor p75, were cultured in DMEM medium (Fisher) supplemented with 10% FBS (Fisher) at 37 °C in a humidified atmosphere with 5% CO2. PC12 and PC12nnr5 cells were cultured in DMEM with 10% FBS and 5% horse serum. Transfection in Hela cells was performed using polyethyleminime (Sigma) at 1-2µg/µl. We found that by using polyethylenimine (PEI) as the transfection reagent in Hela cells the transfection is suboptimal (10-15% of cells transfected) that allow having a small amount of TrkA expressed in the cells and with. As a comparison using the same PEI/DNA ratio in Hek293 cells TrkA is expressed in higher amounts and ligand-independent activation is seen at this quantities of TrkA DNA. 500-1000 ng of DNA per plate was used in TrkA activation experiments. 24h after transfection cells were lifted and re-plated in 12-well plates with 100,000 cells per well. Using this procedure the percentage of transfection is identical in all the wells. 48h after transfection the cells were starved with serum free medium for 2h and stimulated with NGF (Alomone) at the indicated concentrations and time intervals. Cells were lysed with TNE buffer (Tris-HCl pH 7.5, 150 mM NaCl, 1mM EDTA) supplemented with 1% triton X-100 (Sigma), protease inhibitors (Roche), 1 mM PMSF (Sigma), 123 mM sodium orthovanadate (Sigma), and 1 mM sodium fluoride 545 (Sigma). The lysates containing p75 were supplemented with iodoacetamide (Sigma) to avoid post-lysate dimer disulfide formation. Lysates were kept on ice for 10 minutes and centrifuged at 13,000 rpm for 15 minutes on a tabletop centrifuge. The lysates were quantified using a Bradford kit (Pierce) and analyzed by SDS-PAGE or used in immunoprecipitation.

### Western blot analysis

Cells were washed in PBS and lysed in cold lysis buffer (50 mM Tris-HCL [pH7.5], 150 mM NaCl, 1 mM EDTA, 0.1% SDS, 0.1% Triton X-100, 1 mM PMSF, 10 mM NaF, 1 mM Na_2_VO_3_, 10 mM iodoacetamide and protease inhibitor cocktail) at 4°C. Cellular debris was removed by centrifugation at 13,000 g for 15 minutes and protein quantification was performed by Bradford assay. Proteins were resolved in reducing and non-reducing SDS-PAGE gels and membranes were incubated overnight at 4°C with the following antibodies: rabbit polyclonal anti-human p75 intracellular domain (1:1000, Promega); mouse monoclonal anti-HA (1:2000, SIGMA); rabbit polyclonal MBP-probe (1:1000, Santa Cruz); rabbit anti-phosphoTyr674/5 (1:1000, Cell Signaling); rabbit anti-TrkA (1:1000, Millipore). Following incubation with the appropriate secondary antibody, membranes were imaged using enhanced chemiluminescence and autoradiography.

### Electroporation of PC12 and differentiation experiments

The electroporation of the different plasmids was carried out with the Multiporator® (Eppendorf). PC12 cells were gwon with DMEM supplemented with 10% FBS and 5% Horse Serum and antibiotics (gentamycin and penicillin). For elecrtoporation cells were grown to 70-80% confluence on a 10 cm plate and washed with PBS. They were then raised with 3 ml of DMEM medium and centrifuged for 2.5 minutes at 500 rpm. The pellet obtained was resuspended in 3 ml of the hypoosmolar electroporation buffer (KCl 25mM, KH_2_PO_4_ 0.3 mM, K_2_HPO_4_ 0.85 mH, pH 7.2) and a viable counting with trypna blue was carried out. 1 × 10^5^ cells, and a concentration of 5 µg / ml of the plasmid of interest (control, wt or mutant) and a concentration of 5 µg / ml of the plasmid with GFP (Green Fluorescent Protein) were transferred to an electroporation cuvette (2 mm wide and 400 µl in volume (Eppendorf)). After optimizing the transfection parameters, it was determined that the best results were obtained with a pulse of 100 µs at 200V, therefore the electroporation was carried out under these conditions. Finally, the cells were seeded on a 6-well plate with 2 ml of DMEM medium supplemented with 5% horse serum (Gibco). At 24 hours after transfection, the cells were treated with NGF (50 ng/mL) in order to induce the differentitaion of neurites as a function of the plasmid. The length of each neurite was quantified from fluorescence microscopy images uisng the ImageJ software. Three independent electroporation experiments were analyzed and at least 100 neurites per each condition was quantified.

## DATA AVAILABILITY

All the data are contained within the manuscript. Chemical shifts from TrkA-TMD and p75-TMD are deposited in the Biological Magnetic Resonance Data Bank BMRB with accession number 25872 for TrkA-TMD and 19673 for p75-TMD.

## SUPPORTING INFORMATION

This article contains supporting information.

## ACKNOWLEDGMENTS

We thank Dr. M.D. Paul for the acquisition of the LAT-FGFR3 control FRET data set.

## FUNDING INFORMATION

This study was supported by the Spanish Ministry of Economy and Competitiveness (MINECO; project BFU2013-42746-P and SAF2017-84096-R), by the Generalitat Valenciana Prometeo Grant 2018/055 to MV), and by NIH GM068619 (to KH). NMR studies of TRKA-TM and p75-TM were supported by the Russian Science Foundation (grant No# 19-74-30014 to A.S.A).

## CONFLICT OF INTEREST

The authors declare that they have no conflicts of interest with the contents of this article.

